# Differential effects of fenofibrate and fenofibric acid on the regulation of liver endothelial permeability

**DOI:** 10.64898/2026.04.16.718907

**Authors:** Marcin Luty, Dibakar Borah, Karolina Szafrańska, Magdalena Giergiel, Katarzyna Trzos, Peter McCourt, Małgorzata Lekka, Jerzy Kotlinowski, Bartlomiej Zapotoczny

## Abstract

**Background and Aims:** Fenofibrate is widely prescribed for hyperlipidaemia and has been associated with rare but severe cases of drug-induced liver injury (DILI), yet its effects on liver sinusoidal endothelial cells (LSECs) remain to be investigated. LSECs maintain a highly permeable specialized sinusoidal barrier characterized by transcellular pores (fenestrations), regulating the bidirectional transfer of circulating compounds to and from the hepatocytes. As drug-induced alterations in fenestration architecture could influence xenobiotic access to hepatocytes, these changes may modulate pathways associated with DILI. Understanding the effects of fenofibrate on LSEC ultrastructure may therefore provide insights into previously underexplored endothelial contributions to hepatic drug responses.

**Methods:** Both fenofibrate and its active metabolite, fenofibric acid, were evaluated for their effects on LSEC ultrastructure, mechanical properties, and functional markers. Atomic force microscopy (AFM) and scanning electron microscopy (SEM) and were used to quantify fenestration architecture. AFM was additionally used to measure cellular mechanical properties, which were interpreted in the context of fluorescence-based quantification of cytoskeletal organization. Gene expression, viability, and cytotoxicity were assessed using PCR-based and biochemical assays.

**Results:** Fenofibrate reduced fenestration number and porosity at both tested concentration (10, and 25 µM). It also decreased the apparent Young’s modulus of LSECs, accompanied by changes in tubulin and actin architecture, without detectable cytotoxicity. In contrast, treatment with fenofibric acid did not result in significant structural or mechanical effects on LSECs, even at higher concentrations.

**Conclusions:** Together, these data identify LSECs as a drug-responsive hepatic cell type for fenofibrate, suggesting that LSECs could represent an underrecognized contributor to the complex, multifactorial processes underlying DILI. This work provides a framework for evaluating endothelial contributions to fenofibrate-associated liver effects in more complex models.

Fenofibrate reduces LSEC fenestrations and metabolic activity at higher concentrations, while its metabolite, fenofibric acid, does not affect LSEC, regardless of its concentration.

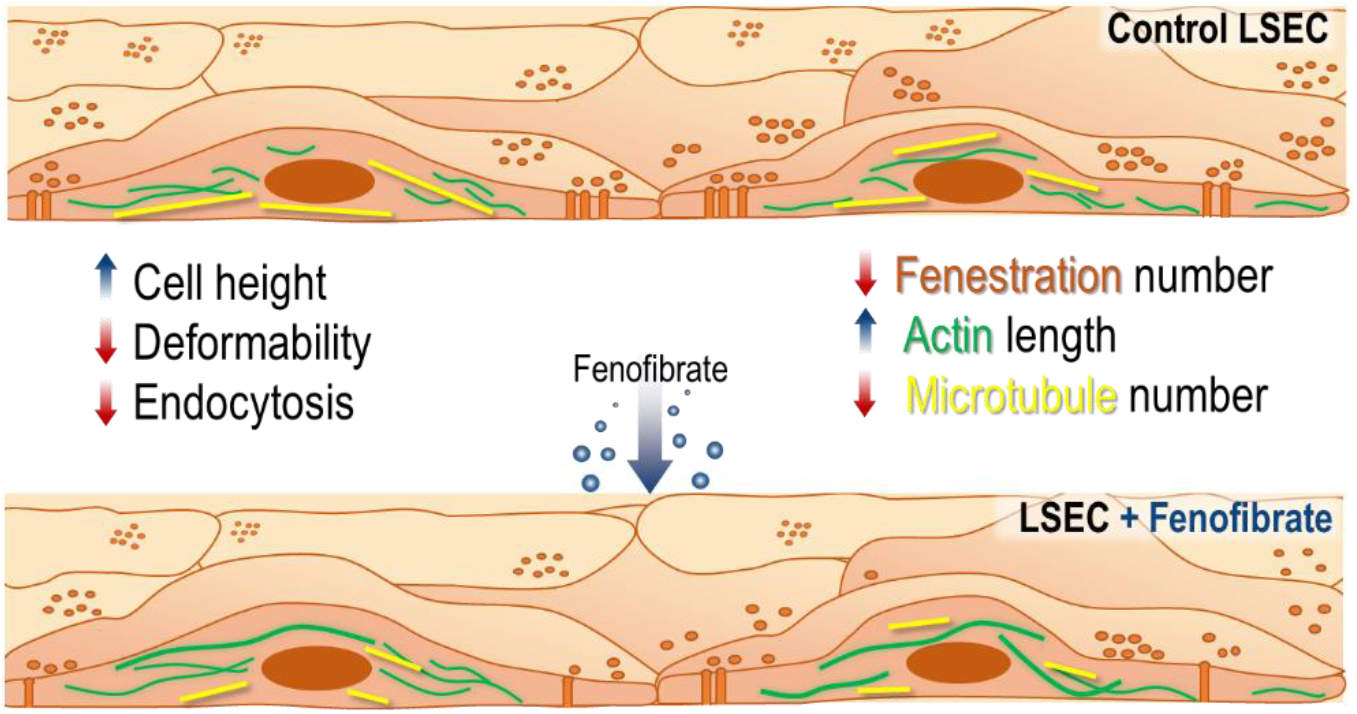

## 1. Introduction

The liver is a crucial organ in detoxifying endogenous and exogenous substances, including pharmaceutical compounds [1]. Consequently, many drugs accumulate in the liver [2,3] and, despite their therapeutic benefits, may cause drug-induced liver injury (DILI) [4,5], a leading cause of acute liver failure [6]. The physical blood-liver barrier in the liver is formed by liver sinusoidal endothelial cells (LSECs). They are specialized endothelial cells characterised by a high endocytic capacity and the presence of fenestrations – transcellular nanopores facilitating the size-selective exchange between the blood and hepatocytes, supporting the clearance of circulating waste products and pathogens [7,8]. Liver dysfunction has been associated with endothelial impairment, particularly with reduced fenestration number in LSECs, which may impair sinusoidal permeability and hepatic drug clearance [9,10].

Fenofibrate (FF) is a drug from the fibrate group that lowers the level of apolipoprotein B and triglycerides in the blood. It also lowers the level of low-density lipoproteins (LDL), while increasing the level of high-density lipoproteins (HDL) [11,12]. FF is a prodrug, converted mainly in the hepatocytes to its biologically active form, fenofibric acid (FFA) [13]. It affects cells by activating peroxisome proliferator-activated receptor alpha (PPARα). These receptors regulate lipid metabolism by activating lipoprotein lipase; they can also stimulate β-oxidation of fats and reduce the production of apolipoprotein C-III, thereby reducing triglyceride production [14,15]. Moreover, FF has been reported to exert broader effects, including anti-inflammatory activity [16], hepatoprotection [17], anticancer properties [18,19], and delayed progression of Duchenne muscular dystrophy [20]. This drug is commonly prescribed to patients with hyperlipidaemia and is frequently administered as a long-term therapy, often for several years [21,22], and can therefore be classified as a chronically used medication. Although FF is generally considered safe in clinical use, potential negative effects of FF on the liver, including the possibility of induction of DILI, are postulated [23,24]. However, the evidence is not uniform, as some studies report hepatoprotective effects of FF in preclinical studies using rodent models [17,25]. Nevertheless, this drug has never before been analyzed on LSECs.

Regarding potential effects on LSECs, FF has been reported to modulate cellular energy metabolism [26], and enhance endothelial barrier function (among other things, by blocking cell diapedesis through them) [27]. Notably, FF has also been demonstrated to reduce reactive oxygen species (ROS) levels [28]. Because elevated ROS contributes to fenestration loss and reduced porosity in LSECs [29], the antioxidant properties of FF suggest a potential role in preserving fenestration dynamics. Our recent studies have demonstrated that ROS promotes fenestration closure, whereas antioxidant treatment with *N*-acetylcysteine or glutathione attenuates these effects [30] and preserves fenestration dynamics [29].

Here, we investigated whether FF and its active metabolite, FFA, influence LSEC morphology, mechanical properties, cytoskeletal organization, and selected gene expression to better define the endothelial role in responses to FF.

## 2. Materials and methods

### Isolation of LSEC

LSECs were isolated from male C57BL/6 mice using the protocol described by *Smedsrød and Pertoft* [31], later modified [32]. Briefly, endothelial-specific CD146 magnetic MicroBeads (MACS, MiltenyiBiotec, Germany) were used for positive immunomagnetic separation of CD146-positive cells (the marker for endothelial cells). Then cells were centrifuged (10 min; 300G) and resuspended in EGM-2 (Lonza) medium at 10^6 cells/ml density at 4°C.

### Cell culture

After isolation, LSECs were seeded onto ø35 mm Petri dishes (65,000 cells/slide; 65 µl of suspension) or coverslips (ø13 mm) coated with fibronectin (40,000 cells/slide; 40 µl of suspension) in a 24-wellplate or 48-well plates (250,000 cells/well 250 µl of suspension); the cell suspension was placed firstly only in the center of the dish or slide or onto the whole surface of well. After 15 minutes of seeding, the dishes were filled with EGM-2 medium. Finally, after 1 h, the medium was replaced with preheated fresh EGM-2 medium containing 0, 10, or 25 µM FF or FFA, and the cells were incubated overnight at 37°C in 5% CO2. Experiments on cells were conducted within 20h after cell isolation.

### MTS assay

To determine cell metabolic activity, the MTS colorimetric assay (Promega, Poland) was performed. In previously prepared cells on 48-well plates, after 16 h of incubation with FF, the medium was replaced with 300 µl of fresh EGM-2 (containing FF at the same concentration) containing tetrazolium salts. After 2 h, 100 µl of medium was collected from each well on a 96-well plate, and then the absorbance was measured at 492 nm using a LEDETECT 96 microplate reader (Labexim Products, Austria).

### LDH Assay

To determine FF toxicity, we used the CyQUANT™ LDH Cytotoxicity Assay Kit from Invitrogen. This assay detects lactate dehydrogenase (LDH), which is released into the medium upon cellular injury.

Cells were plated in a 48-well plate. After 16 hours of cell incubation in the presence of FF, 50 µl of culture medium was collected in a 96-well plate and 50 µl of assay reagent was added to it, and after 30 minutes, the reaction stop reagent (30 µl) was added. Absorbance was measured at 492 nm using a LEDETECT 96 microplate reader (Labexim Products, Austria). FF toxicity was determined relative to the absorbance obtained from lysed cells, indicating the release of all LDH from all cells.

### Analyses of cell elasticity

Cells were seeded on the surface of plastic Petri dishes as described (cell culture). Before AFM measurements, the medium was removed, and cells were washed twice with 37°C EGM2. The medium was replaced with 1.1 ml of CO_2_-independent medium (Gibco, supplemented with an appropriate concentration of FF or FFA). The measurements were conducted using atomic force microscopy (AFM, Nanowizard 4 head, Bruker/JPK Instruments) at 37°C, maintained by a PetriDish heater™ (Bruker/JPK Instruments). V-shaped cantilevers with the nominal spring constant of 0.01 N/m were used (MLCT-BIO-DC-c, Bruker). The calibration of the cantilever was conducted on a Petri dish surface without cells. For each condition, at least 10 cells were measured. For each cell, 64 force-distance curves were acquired (loading force: 2 nN, loading rate: 8 µm/s, z range: 4 µm, scan size 8×8 μm). Each sample was measured within 30-50 minutes. Each experiment was performed in triplicate. The analyses of elasticity were conducted using the JPKSPM Data Processing program for cone cantilevers.

### Quantitative imaging

Analysis of the number of fenestrations was performed using atomic force microscopy (AFM) on fixed cells on a Petri dish. After 16 hours of incubation with FF or FFA, cells were washed with PBS at 37°C, then fixed in 2% warm glutaraldehyde solution for 3 minutes. Cells were then washed with PBS solution 3 times. Before analyses, cells were washed twice with PBS and suspended in PBS with penicillin and streptomycin (Merck). Analyses were performed using an AFM (Nanowizard 4 head, Bruker/JPK Instruments) at 25 °C on a PetriDish heater™ (Bruker/JPK Instruments). Measurements were performed in Quantitative Imaging (QI) mode, using SMC-PIC-V2 cantilevers with a nominal spring constant of 0.1 N/m. Parameters were adjusted to the scan frame and required resolution, but were usually in a range of: loading force: 0.2-0.4 nN, loading rate: 100-135 nm/s, z range: 950-1300 nm, scan size: 30×35 µm^2^ or larger, resolution: >12 pixels/µm. The AFM images were post-processed using the JPKSPM Data Processing software and prepared for further analysis.

### Analyses of fenestrations

Within the selected membrane areas of at least 1000 µm^2^, excluding the nuclear area and glass surface, fenestrations were identified automatically using a previously described neural-network-based analysis of AFM force-spectroscopy maps [33]. The classification relied on two physical parameters reflecting local membrane properties: relative cell surface height, which captures pore topography, and local stiffness, which provides complementary information on the mechanical properties of the membrane and the underlying substrate. By combining these two features, the method recognizes fenestrae within sieve plates and quantifies the number of fenestrations and their approximate sizes across the analyzed membrane regions.

### Staining of the cells

LSECs were grown on coverslips, after which cells were fixed (3.7% formaldehyde; 20 min) and permeabilized (0.2% Triton-X100; 7 min). After each step, the cells were washed 3 times with PBS. Then the cells were incubated overnight in the presence of primary antibodies (Merck anti-tubulin mouse antibodies 1:200). Cells were then incubated with secondary antibodies (Alexa 594™ goat anti-mouse antibody; 1:100 dilution in PBS, Invitrogen, A11005, excitation/emission 590/618 nm; 2 hours incubation). Actin filaments were labeled by incubating cells with phalloidin solution (Alexa Fluor 488™ 1:200 in PBS, Invitrogen, A12379, excitation/emission 495/518 nm; 60 min). Cell nuclei were labeled by incubating them with Hoechst 34 580 solution (1:5000 dilution in PBS, Molecular Probes, H21486, excitation/emission 392⁄440; 15 min). The cell slides were mounted to microscope slides using mounting medium (Prolong Diamond)

### Endocytosis

The LSECs were seeded in fibronectin-coated 48-well plastic plates (300000 cells per well). For the endocytosis experiments, the cell media were supplemented with 1% human serum albumin, 30 ng/mL of radiolabeled formaldehyde-treated serum albumin (125I-FSA), and various concentrations of unlabeled FSA (0/30/75/150 µg/mL) and incubated for 2 h. Then, the cell media were removed and undigested ligands were precipitated with 20% trichloroacetic acid to calculate the degradation level [34]. To calculate the number of cells in each well, the cell nuclei were stained with Hoechst 33258 (1:1000) for 20 min at 37 °C. Images obtained using a fluorescence microscope (EVOS, ThermoFisher) were analyzed using simple threshold-based segmentation in Fiji [34]. Next, cells were lysed with 1% sodium dodecyl sulfate (SDS) to analyze the cell-associated fraction of FSA. All experiments were performed with 3 bioreplicates/animals and 3 technical replicates per experiment. Total endocytosis (both the cell-associated and degraded fractions) was normalized to the calculated number of cells in each well.

### SEM

Isolated LSEC were seeded on fibronectin-coated 16-well plates (CS16-CultureWell™ Removable Chambered Coverglass, Grace Bio-labs and after treatments with 0, 10, and 25 μM FF in EGM-2 for 16h fixed in 4% formaldehyde and 1% glutaraldehyde in PHEM (pH 7.2) and stored in fixative until further processing. Samples were then washed in PHEM before 1 h incubation with 1% tannic acid (freshly prepared in PHEM) followed by 1 h in 1% osmium tetroxide in H2O and dehydration in an ethanol gradient (30%, 60%, 90%, 4×100% for 5 min each) and chemical drying in hexamethyldisilane twice for 2 min (Sigma-Aldrich, Oslo, Norway). Before imaging, samples were mounted on aluminum stubs and coated with a 10 nm layer of gold/palladium alloys.

Images of randomly selected 35-60 cells per sample, from four separate areas, were taken using SEM Gemini 300 (Zeiss) at 2kV with pixel size 25 nm. All treatments were performed on 3 separate biological replicates. Fenestrations were analyzed using an automated segmentation tool based on StarDist [35] as described in detail in [36]. Segmentation provided quantitative data including fenestration frequency (number of fenestrations per cell area µm^2^), porosity (% of fenestration-occupied cell area) and diameter of the fenestration (assuming circularity).

### Cytoskeleton visualization

Actin filaments and microtubules of previously stained cells (cell staining) were visualized using a fluorescence microscope (Olympus IX83 inverted microscope with a 100 W mercury lamp and Photometrics Prime BSI Express camera).

### Cytoskeleton analyses

The numerical analyses of the actin cytoskeleton and microtubules were performed using the filament sensor program [37]. Cell images obtained with an Olympus fluorescence microscope were processed in ImageJ (Fiji) to isolate single cells. The images prepared in this way were uploaded together (all for a given condition) to the filament sensor program. The images were analyzed without pre-processing. Only filaments longer than 5 µm (75 pixels) for Actin and 3,5 µm for Tubulin were taken into analysis.

### RNA isolation and real-time PCR analyses

Total RNA from LSEC was isolated using Fenozol (A&A Biotechnology). Next, 500 ng of RNA was used for reverse transcription using oligo(dT) primers (Promega) and M-MLV reverse transcriptase (Promega). Quantitative PCR (qPCR) was carried out using SYBR Green Master Mix (Sigma-Aldrich) and a QuantStudio qPCR System (Applied Biosystems). Gene expression was normalized to Ef2, and then, the relative transcript level was quantified by the 2^deltaCt method. Primer sequences (Genomed/Sigma-Aldrich) are listed in Table 1.

**Supplementary Table 1.**
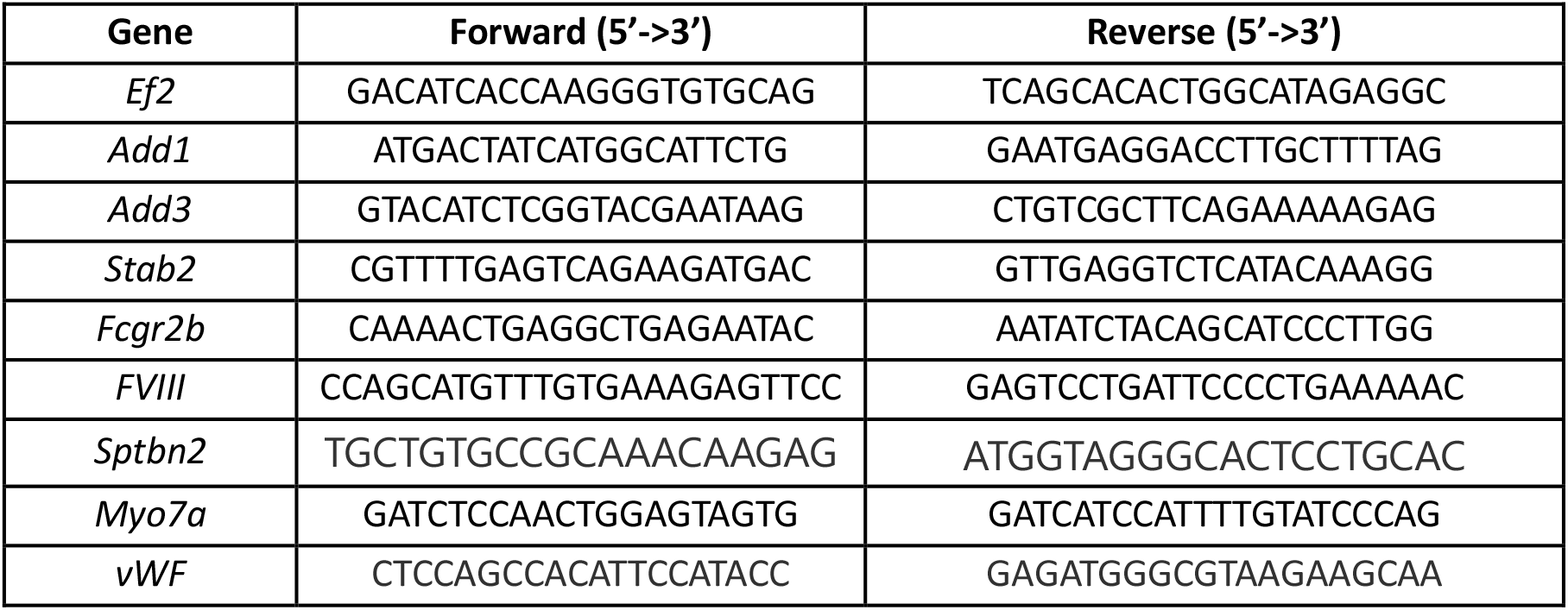
Sequences of primers used for qPCR.

### Statistical analysis

All data were checked for normality of distribution using the Shapiro-Wilk test. Normally distributed data were compared using the ANOVA test with the post-hoc Tukey test. For nonparametrically distributed data, the comparison was performed using the Kruskal-Wallis test with the post-hoc Dunn’s test. On graphs, the differences are presented with * for p<0.05; ** for p<0.01 and ^***^ for p<0,001.

## 3. Results

To assess cytotoxic effect of FF and FFA, cell metabolic activity was measured by the MTS assay, and membrane integrity was evaluated by LDH release. MTS assays showed that FF treatment reduced mitochondrial activity of LSECs by two-fold (**Fig. 1A**). LDH release was not significantly altered, indicating nondetectable membrane damage after the treatment (**Fig. 1B**). FFA treatment did not show any marks of cytotoxicity or changes in membrane integrity (**Fig. S1**).

**Fig. 1.**
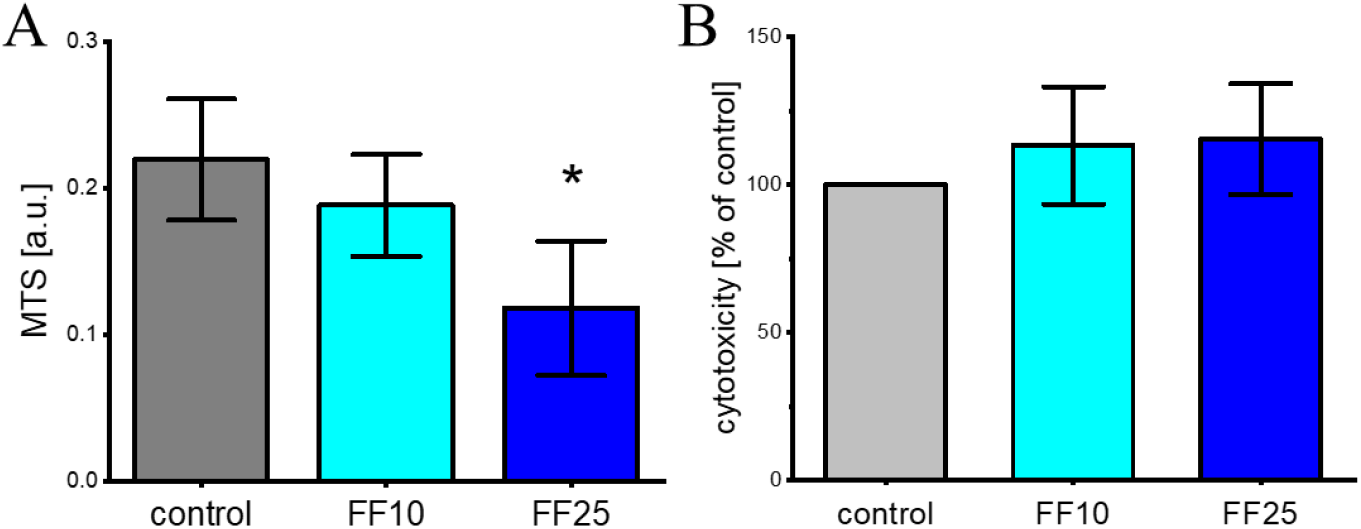
Cytotoxic effect of FF. A) Mitochondrial activity of LSECs after 16h incubation with FF, assessed by the MTS assay; B) LDH release (term as cytotoxicity; the value is standardized relative to the control) after 16h incubation with FF, Columns represent mean values; whiskers indicate standard deviation (SD). Statistical significance was assessed using the Kruskal-Wallis test with Dunn’s post hoc test; * *p* < 0.05,; n = 3.

LSEC fenestrations are critical for the exchange of solutes, lipoproteins, and small particles between blood and hepatocytes. Since fenofibrate can modulate lipid metabolism and cytoskeletal dynamics, we investigated whether it alters LSEC morphology. QI mode AFM analysis showed that fenofibrate induced a dose-dependent reduction in LSEC porosity, with the strongest effect observed at 25 µM (**Fig. 2A, B**). Topography analysis further revealed that FF treatment at either 10 µM or 25 µM increased LSEC cell height by approximately 20% (**Fig. 2C**). In contrast, FFA did not induce defenestration and did not affect LSEC cell height at either concentration (**Fig. S2**).

**Fig. 2.**
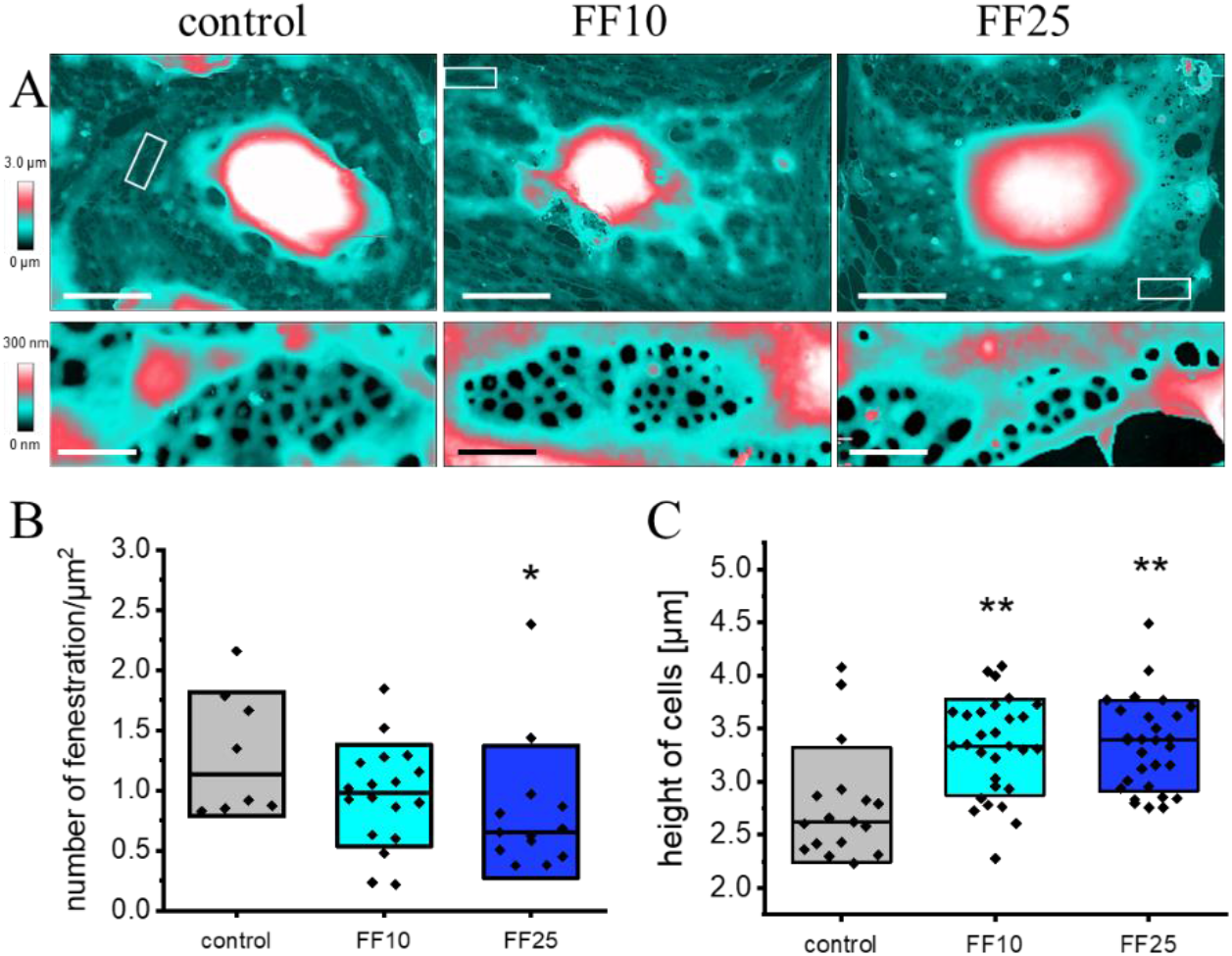
The effect of FF on LSEC fenestrations measured using AFM. A) Representative AFM topography images of LSECs treated with 0, 10, and 25 μM FF (scale bar = 10 μm). The white rectangle marks the region enlarged in the bottom row, highlighting sieve plates with fenestrations (scale bar = 1.0 μm). B) Fenestration frequency (number of fenestrations per μm^2^). C) Cell height, defined as the maximum height measured in the central region of the cell. *Boxes represent SD, dots represent individual cells*. Statistical significance was assessed using the Kruskal-Wallis test with Dunn’s post hoc test; *p* < 0.05, \**p* < 0.01, \*\**p* < 0.001; n = 4.

SEM was used for high-throughput evaluation of the effects of the FF treatment on LSEC morphology (**Fig. 3**). FF treatment shifted fenestration size distribution toward larger diameters (**Fig. 3 B, C**). Compared to control, both 10 and 25 µM FF reduced the proportion of fenestrations in the 100-200 nm range (from 84.69% to 80.12% and 77.08%, respectively) and increased the fraction of fenestrations > 200 nm (from 9.23% to 12.58% and 12.77% for 10 and 25 µM FF, respectively). An increase in < 100 nm fenestrations was also observed, more pronounced at the higher dose (**Fig. 2C**). The data indicate a dose-dependent enlargement and broadening of fenestration size distribution. The division of fenestrations into such groups is related to the fact that fenestrations of 100-200 nm in size are those that function most optimally [34]. Similar to our observations in AFM, SEM data show that changes in fenestration size did not compensate for the reduction in fenestration number (**Fig. 3D**; fenestration frequency), resulting in a net decrease in porosity, expressed as the percentage of cell surface area occupied by fenestrations (**Fig. 3E**).

**Fig. 3.**
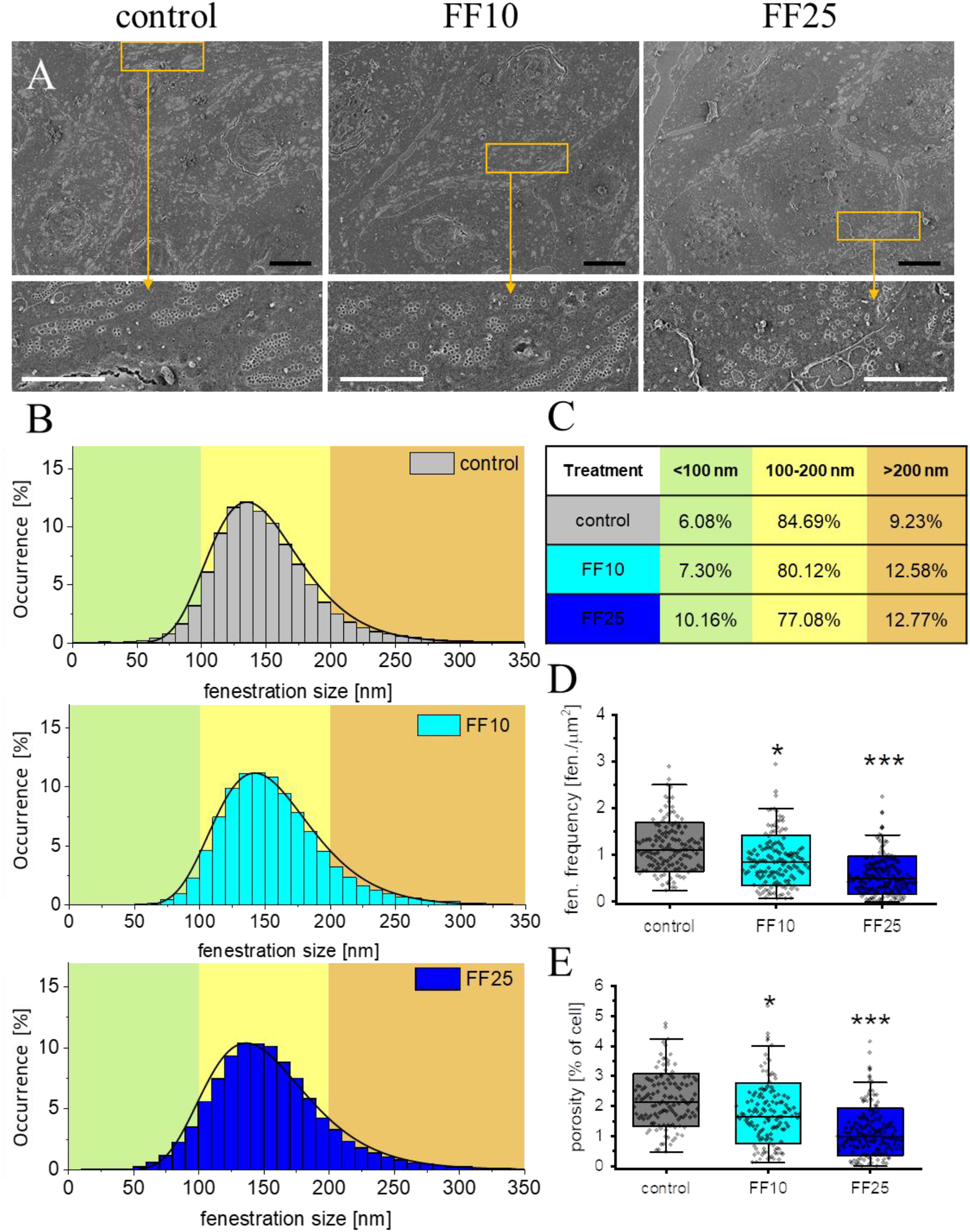
The effect of FF on LSEC fenestrations measured by SEM. A) Representative SEM images of LSECs treated with 0, 10, and 25 μM FF (scale bar = 10 μm). The yellow rectangle in the pictures in upper row marks the region enlarged in the bottom row, highlighting sieve plates with fenestrations (scale bar = 2.0 μm). B) Fenestration diameter distribution Black line indicates for a log-normal function fit to the data. Over 100,000 fenestrations were measured per condition. C) the table contains ratio of fenestration size within three predefined diameter ranges: small (<100 nm, green), intermediate (100-200 nm, yellow) and large (>200 nm, orange). D) Fenestration frequency described as the number of fenestrations per area. E) Porosity defined as the fraction of cell surface covered by fenestrations. Boxes in D) and E) represent SD; dots refer to individual fenestrations; black line within the box indicates median and whiskers are 5th and 95th percentiles of the data. Statistical significance (ANOVA with post-hoc Dunnett’s test): \**p* < 0.05, \*\**p* < 0.01, ^***^*p* < 0.001; n = 3.

To investigate whether cytoskeletal remodeling contributes to FF-induced (and FFA-induced) changes in LSEC fenestration, actin filaments and microtubules were visualized and quantitatively analyzed using fluorescence microscopy (**Fig. 4, Figs. S3-S4**). Treatment with 25 µM FF resulted in more than twofold reduction in the number of long (> 5 µm) actin filaments (**Fig. 4B**) accompanied by a modest increase in filament length (**Fig. 4C**). In contrast, FF significantly increased the number of microtubules (**Fig. 4D**). The increase in number was the most pronounced for short filaments (3.5-5.0 µm; **Fig. S4**) and was accompanied by a reduction in their average length (**Fig. 4E**). Additional experiments with FFA showed no statistically significant alterations in actin filament or microtubules organization in LSECs (**Fig. S5**).

**Fig. 4.**
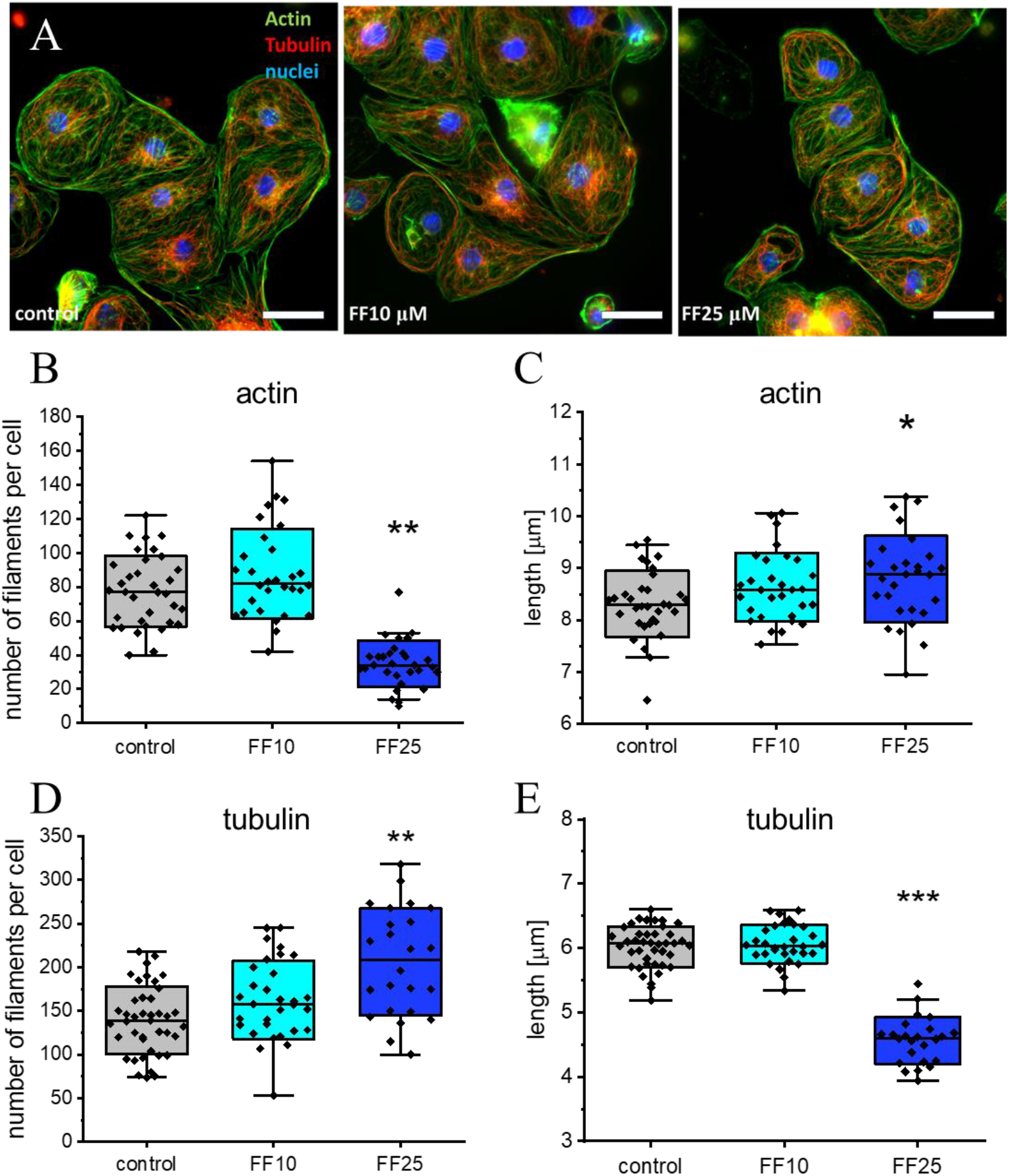
Quantitative fluorescence microscopy of actin and tubulin filaments. A) representative images of LSECs treated with 0, 10 and 25 μM FF. Actin (green) tubulin (red), cell nuclei (blue); scale bar = 25 µm; B) Number of actin filaments per cell longer than 5.0 μm. C) Average length of actin filaments in a cell (for filaments >5 μm); D) number of microtubules > 3.5 μm per cell; E) average length of microtubules per cell (for filaments >3.5 μm). Boxes in B-E represent SD; dots refer to individual cells; square symbol indicates median and whiskers are 5th and 95th percentiles of the data. Statistical significance was assessed using the Kruskal-Wallis test with Dunn’s post hoc test; *p* < 0.05, \**p* < 0.01, \*\**p* < 0.001; n = 3.

Changes in the cytoskeleton are often accompanied by alterations in cell elasticity, which can be quantified as the apparent Young’s modulus using AFM. Accordingly, the elasticity of LSECs exposed to FF was evaluated (**Fig. 5**). The apparent Young’s modulus was 9.6 ± 5.9 kPa in the control group and decreased to 5.0 ± 3.5 kPa and 5.2 ± 3.9 kPa following treatment with 10 and 25 µM FF, respectively (**Fig.5**. Both concentrations produced a comparable effect, resulting in an approximately twofold reduction in cell stiffness. In comparison, FFA induced only a minor, statistically insignificant reduction in the apparent Young’s modulus of LSECs (**Fig. S6**), reflecting observed minor changes in the architecture of the cytoskeleton.

**Fig. 5.**
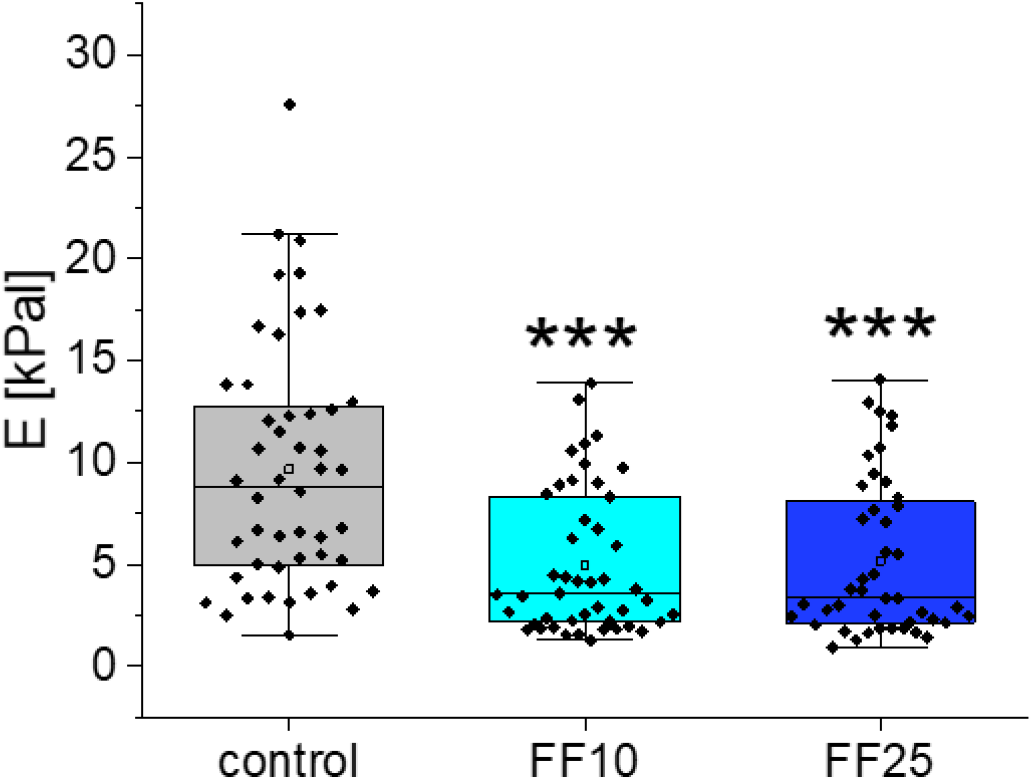
Apparent Young’s modulus of LSECs after FF treatment. Apparent Young’s modulus determined from AFM force-indentation curves at a defined indentation depth. Values are reduced for 10 and 25 µM FF compared to control. Boxes indicate the standard deviation; dots represent the mean apparent Young’s modulus obtained from force-indentation maps for individual cells, whiskers are 5th and 95th percentiles of the data. Statistical significance was assessed using the Kruskal-Wallis test with Dunn’s post hoc test; *p* < 0.05, \**p* < 0.01, \*\**p* < 0.001; n = 3.

The effect of FF on LSEC endocytic capacity was assessed using an FSA-based assay. While 10 µM FF did not affect endocytic function, treatment with 25 µM FF reduced the cell-associated fraction of FSA (**Fig. 6A**). The fraction of degraded FSA remained unchanged, indicating that intracellular degradation was not affected. Together, these data suggest that FF at higher concentrations potentially alters receptor-mediated uptake without impairing lysosomal degradation.

**Fig. 6.**
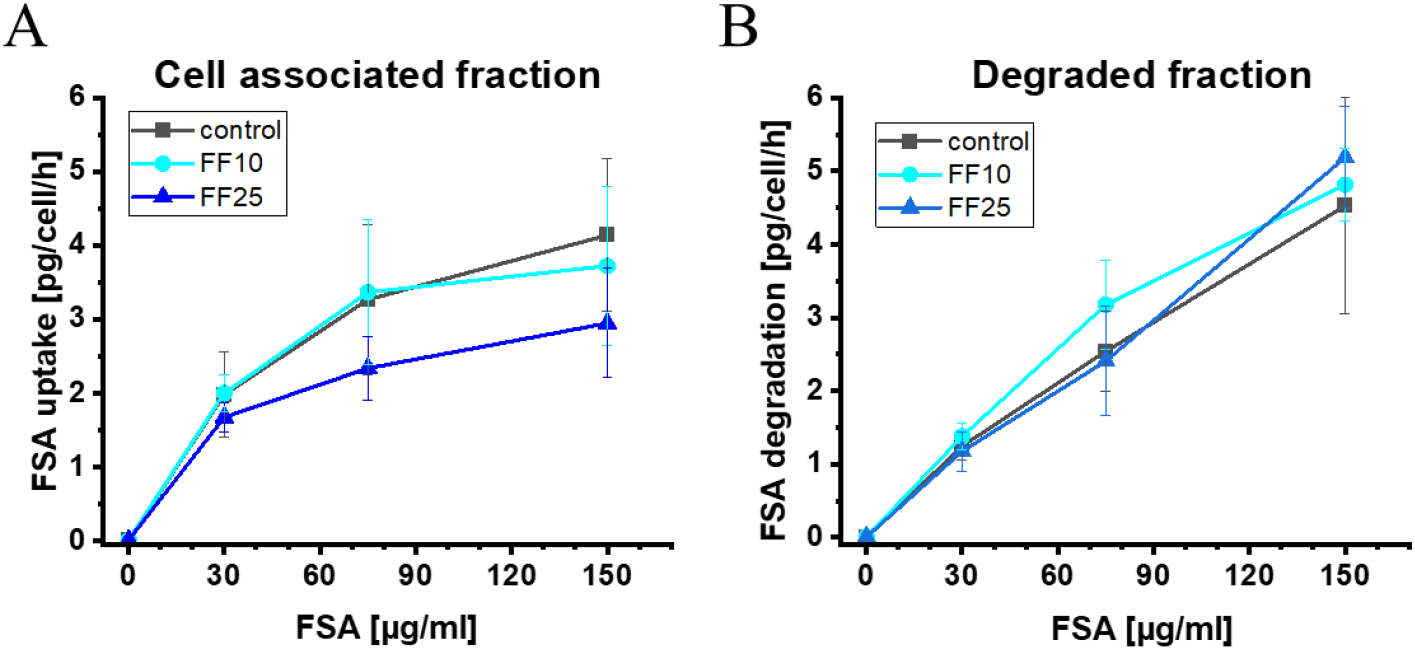
Effect of FF on LSEC scavenging function. Scavenger receptor-mediated endocytosis of ^125^I-FSA measured after 0, 10 and 25 µM FF treatment with increasing dose of FSA. A) Fraction of FSA ligand associated with cells; B) degraded fraction of FSA. Whiskers represent standard deviation from n = 3 bioreplicates.

To assess whether FF-induced structural changes involve transcriptional regulation, mRNA levels of genes associated with LSEC function (*FVIII, Stab2, vWF2, Fcgr2b*) and fenestration architecture (*Add1, Add3, Sptbn2, Myo7a*) were analyzed. 25 µM FF treatment reduced expression of *Myo7a, Stab2* and *Fcgr2b* (**Fig. 7A, B**).

**Fig. 7.**
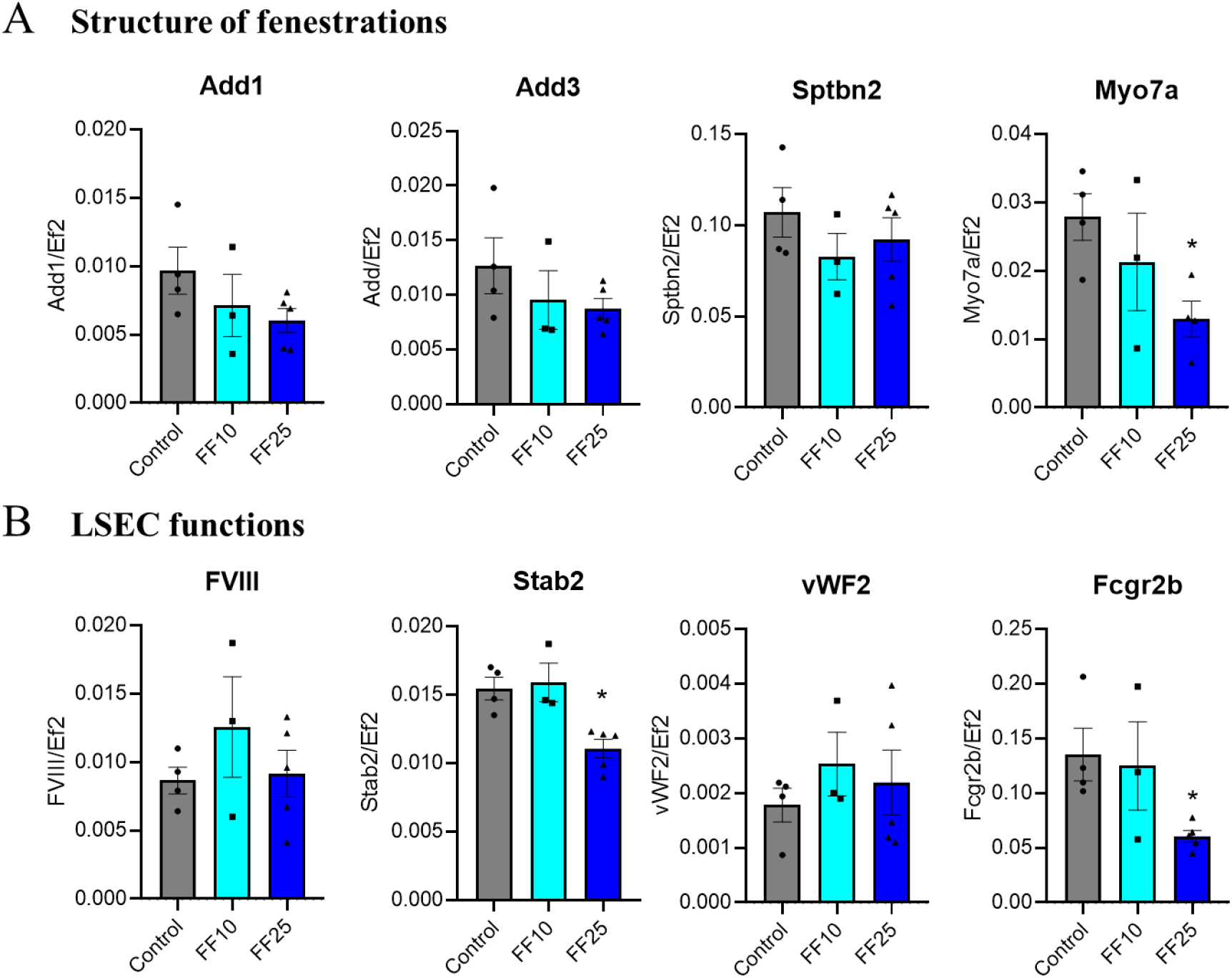
Relative mRNA levels of genes associated with fenestration architecture (A) and LSEC function (B). Columns represent mean values; points represent biological replicates (n = 3-6); whiskers indicate standard deviation. Statistical significance was assessed using the Kruskal-Wallis test with Dunn’s post hoc test; *p* < 0.05, \**p* < 0.01, \*\**p* < 0.001.

## 4. Discussion

This study demonstrates that fenofibrate (FF) exerts concentration-dependent effects on the structure and mechanical properties of liver sinusoidal endothelial cells (LSECs). In particular, higher concentrations of FF reduced fenestration number, altered cytoskeletal organization, and decreased apparent Young’s modulus, whereas its active metabolite, fenofibric acid (FFA), did not produce comparable changes under the conditions examined [13,38,39]. Both compounds were included in this study because, following oral administration, FF is delivered to the liver via the portal vein, where LSECs represent one of the first cellular barriers exposed to the drug. Although FF is rapidly hydrolyzed to FFA by hepatic carboxylesterases [13], transient exposure of LSECs to the parent compound during first-pass metabolism is likely. Given the central role of LSECs in regulating sinusoidal permeability and hepatic microenvironment homeostasis [7,10], such exposure may be sufficient to induce structural and functional alterations in LSEC which could influence hepatic responses to xenobiotics and contribute to processes associated with DILI.

Key aspects of sinusoidal permeability facilitating the exchange of circulating compounds between blood and hepatocytes are the presence of fenestrations and high cellular endocytosis of LSECs. Both ensure selective and bidirectional transport which is fundamental for liver function [8,40,41]. Observed here, the reduction in the number of fenestrations and reduced cell-associated fraction of FSA may contribute to decreased LSEC clearing efficiency. In particular, Stabilin-2 is a major receptor responsible for the uptake of oxidized and glycosylated ligands, whereas Fcgr2b plays a critical role in immune complex internalization and regulation of inflammatory responses [8, 42,43]. The latter is not only connected with immune function [44] but also critical for endocytosis in cells [45] including LSECs [43]. The observed downregulation of Stabilin-2, at higher FF concentrations is therefore consistent with the reduced endocytic capacity of FSA which is primarily endocytosed by stabilin-1 and stabilin-2 [46]. Importantly, these changes occurred in the absence of detectable cytotoxicity, indicating that LSEC filtration impairment represents a specific structural-functional response rather than a consequence of cell death.

The results presented in this study showing a decrease in LSEC stiffness after FF could indicate that this drug improves cellular function. Previous studies have shown that reductions in apparent Young’s modulus were associated with actin depolymerization and reorganization into mesh-like structures that may facilitate fenestration formation [8,32]. Proteins such as spectrin play an important role in this process by linking the actin cytoskeleton to membrane [47]. However, cytoskeletal regulation of fenestrations is complex; for example, inhibition of protein disulfide isomerase (PDI) by Bepristat altered fenestration number without affecting nanomechanical properties, indicating that structural and mechanical parameters are not always directly coupled [48]. In the present study, quantitative fluorescence microscopy revealed changes in both actin and microtubule organization at higher FF concentrations, while reduced cellular stiffness was equally significant for 10 and 25 µM FF. These findings indicate that more detailed changes in the cytoskeletal architecture are present but cannot be resolved by diffraction-limited microscopy and require developing more advanced approaches. Another possibility is that the reduction in stiffness is mediated in part by cellular relaxation through inhibition of actomyosin contractility [49,50]. Here, we observed a decreased expression of *Myo7a*, encoding myosin VII, which was reported as an important regulator of fenestration size through interactions with the actin cytoskeleton [32,51,52]. In line with previous work demonstrating that myosin-dependent mechanisms regulate fenestration size and distribution (Rho–ROCK signaling; [32,53]), the observed downregulation of *Myo7a* may contribute to the observed broadening of fenestration size distribution and increased fraction of large fenestrations after FF. In summary, cellular nanomechanics represents an important, yet not fully understood, determinant of LSEC structure and function.

Quantitative analysis of microtubules showed an increase in their abundance following FF treatment, which could partially compensate for the reduction in cellular stiffness [54]. Microtubules also play a critical role in endocytosis and intracellular trafficking [55,56], yet their increased abundance did not translate into enhanced endocytic activity. Instead, higher concentrations of FF were associated with a reduction in endocytic capacity. These counterintuitive findings can be explained by the fact that FF is a highly lipophilic compound with low aqueous solubility, properties that favor its accumulation within cellular and mitochondrial membranes. Such accumulation has been associated with alterations in mitochondrial function, including inhibition of respiratory chain activity and changes in cellular energy metabolism [57,58]. These effects may influence cellular processes beyond observed biomechanical changes, including reduced endocytic function.

This study shows that fenofibrate has a dose-dependent effect, with 25 μM potentially impairing LSEC function in liver sinusoids. It should be emphasized that the FF concentrations used in the analyses represent its biologically available concentrations [59] and were selected based on pharmacokinetic data of the concentration of this drug in the plasma of patients taking FF [21]. Owing to its high lipophilicity and low water solubility, fenofibrate can accumulate in cellular and mitochondrial membranes, which may contribute to defenestration and partially inhibit the mitochondrial electron transport chain, further affecting cellular function [57,58]. The distinct effects of FF and FFA may be related to their physicochemical and pharmacokinetic properties. FFA exhibits greater hydrophilicity and improved bioavailability compared to FF [60,61], which may result in different patterns of cellular exposure and reduced membrane accumulation. In contrast, the higher lipophilicity of FF may promote its accumulation in cellular membranes, potentially contributing to the structural and functional alterations observed in this study. Nevertheless, reports of adverse *in vivo* effects associated with FFA have also been described [62], indicating that its impact may depend on the biological context.

When considering complications, including DILI reported in some studies following FF or FFA administration [23,62], it should be noted that these drugs are commonly prescribed to patients with metabolic disorders, including obesity and dyslipidaemia, conditions that themselves may lead to liver pathology [63]. Such pathological livers are often associated with impaired LSEC morphofunction, including defenestration [40,48]. It should also be added that with the age of the patient, the number of fenestrations in LSEC decreases [51], and FF is used mainly by elderly people. Therefore, these drugs are administered in a context in which the sinusoidal barrier may already be altered. The negative effects of FF on LSECs observed in this study may contribute to the mechanisms underlying DILI associated with FF treatment [23,62]. Further studies on LSECs in models of liver pathology are required.

## Conclusion

These findings demonstrate differential responses of LSECs to fenofibrate and its active metabolite and point to a functional interplay between cytoskeletal organization, cellular mechanics, and fenestration architecture in regulating endothelial function. The observed alterations indicate that LSECs are sensitive to fenofibrate exposure even in the absence of overt cytotoxicity, suggesting that endothelial responses may represent a previously underrecognized component of fenofibrate-associated liver effects. Further studies in more complex and *in vivo* models are required to determine the relevance of these findings for drug-induced liver injury.

## Supporting information

https://1drv.ms/f/c/e91d2347465fb17a/IgBZDTaMUcKdTaEw7DlhNnTjAWsuF8jYJJudvwM2nqpNSwo?e=VfS0IF

## Acknowledgements

The authors thank Jakub Pospíšil (UiT The Arctic University of Norway, Tromsø) for support with automated fenestration analysis in SEM micrographs. The authors also thank Joanna Pabijan (Institute of Nuclear Physics, Polish Academy of Sciences) for continuous support in the biological laboratory.

