## Supplementary material for "Differential effects of fenofibrate and fenofibric acid on the regulation of liver endothelial permeability": https://1drv.ms/f/c/e91d2347465fb17a/IgBZDTaMUcKdTaEw7DlhNnTjAWsuF8jYJJudvwM2nqpNSwo?e=VfS0IF

* Authors to whom correspondence

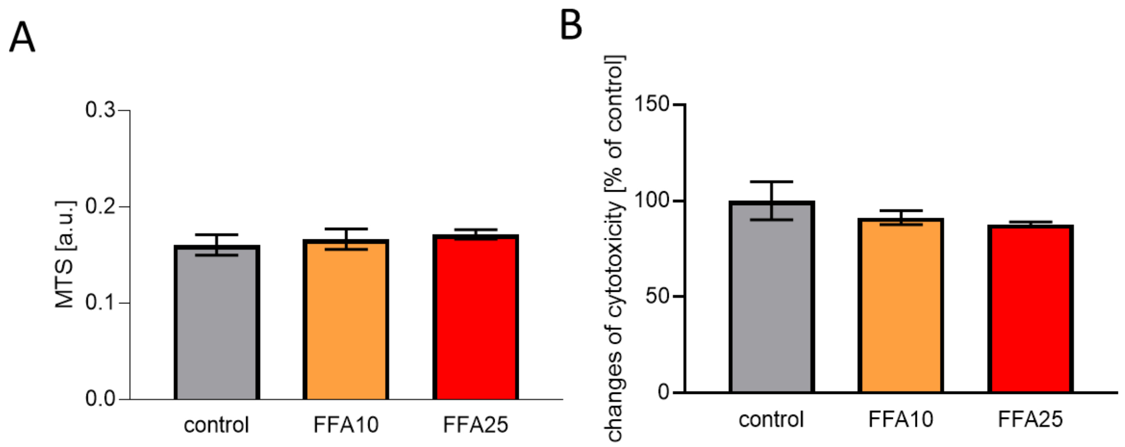


**Supplementary information 1 Cytotoxic effect of fenofibric acid** A) Mitochondrial activity (MTS assay) of LSECs after 16h incubation with FFA; B) LDH assay. Columns represent mean values; whiskers indicate standard deviation (SD). Statistical significance (Kruskal-Wallis test with the post-hoc Dunns test * $p < 0.05$, ** $p < 0.01$, *** $p < 0.001$. n = 3

FF is metabolized to fenofibratic acid (FFA), so we also conducted analyses of the effect of FFA on LESCs. These studies showed that FFA did not exhibit cytotoxic effects on LSECs, but it also did not reduce their metabolic activity (fig. S1), in contrary to FF (fig. 1).

Changes in the number of fenestrations are a marker of LSEC well-being, so we conducted analyses of the effect of FFA on this aspect. These analyses showed that this drug did not lead to statistically significant changes in the number of fenestrations in LESCs (fig. S2), as it was observed in the case of FF (fig. 2).

Additionally, we analyzed the effect of FFA on the actin cytoskeleton of LESCs using immunochemical staining (fig. S5) and subsequent analysis of these staining using the filament sensor 2.0 program. This program automatically detects actin filaments, but also microtubules, determining their number, length, and thickness (fig. S4). These analyzes showed that FFA did not induce changes in either the number or length of actin filaments (fig. S4) in contrary to FF which was reducing the number of actin filament especially the smaller ones (5-8 µm fig. 5). Moreover, cell elasticity analyzes showed that FFA does not induce changes in the yang module in LSECs (fig. S6).

These results show that FFA is even safer to use than FF, which at higher concentrations led to some disturbances in the functioning of LSEC.

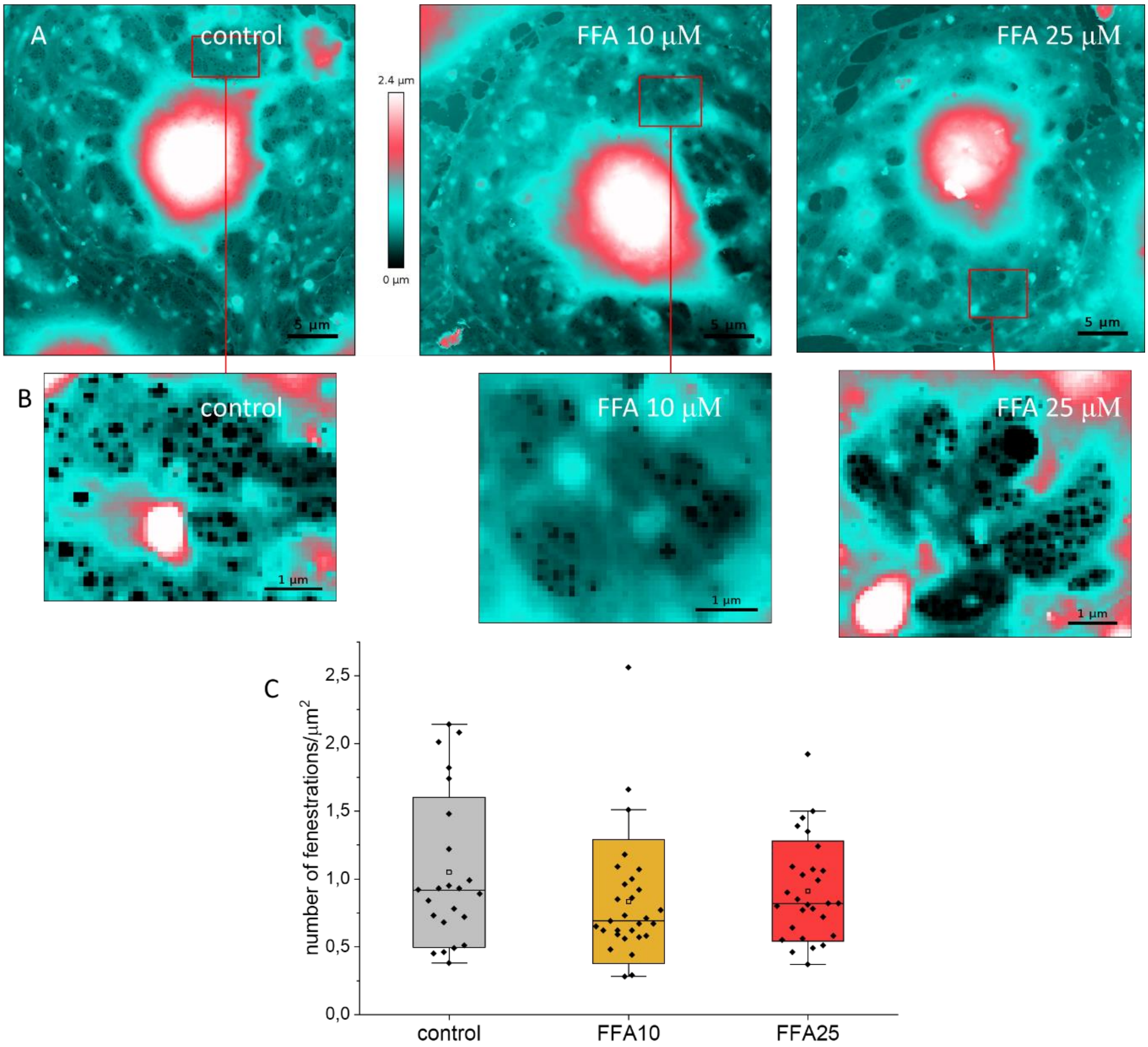


**Supplementary information 2 The effect of FFA on LSEC fenestrations**. A) QI scans of cells after 16 h of incubation in the presence of the drug; (bars represent 5 um length); B) QI scans of sieve plates (bars represent 1 µm length) C) number of fenestrations per µm$^2$. *Boxes represent standard deviation; dots represent individual cells. Statistical significance (Kruskal-Wallis test with the post-hoc Dunns test): * $p < 0.05$, ** $p < 0.01$, *** $p < 0.001$. n =4.*

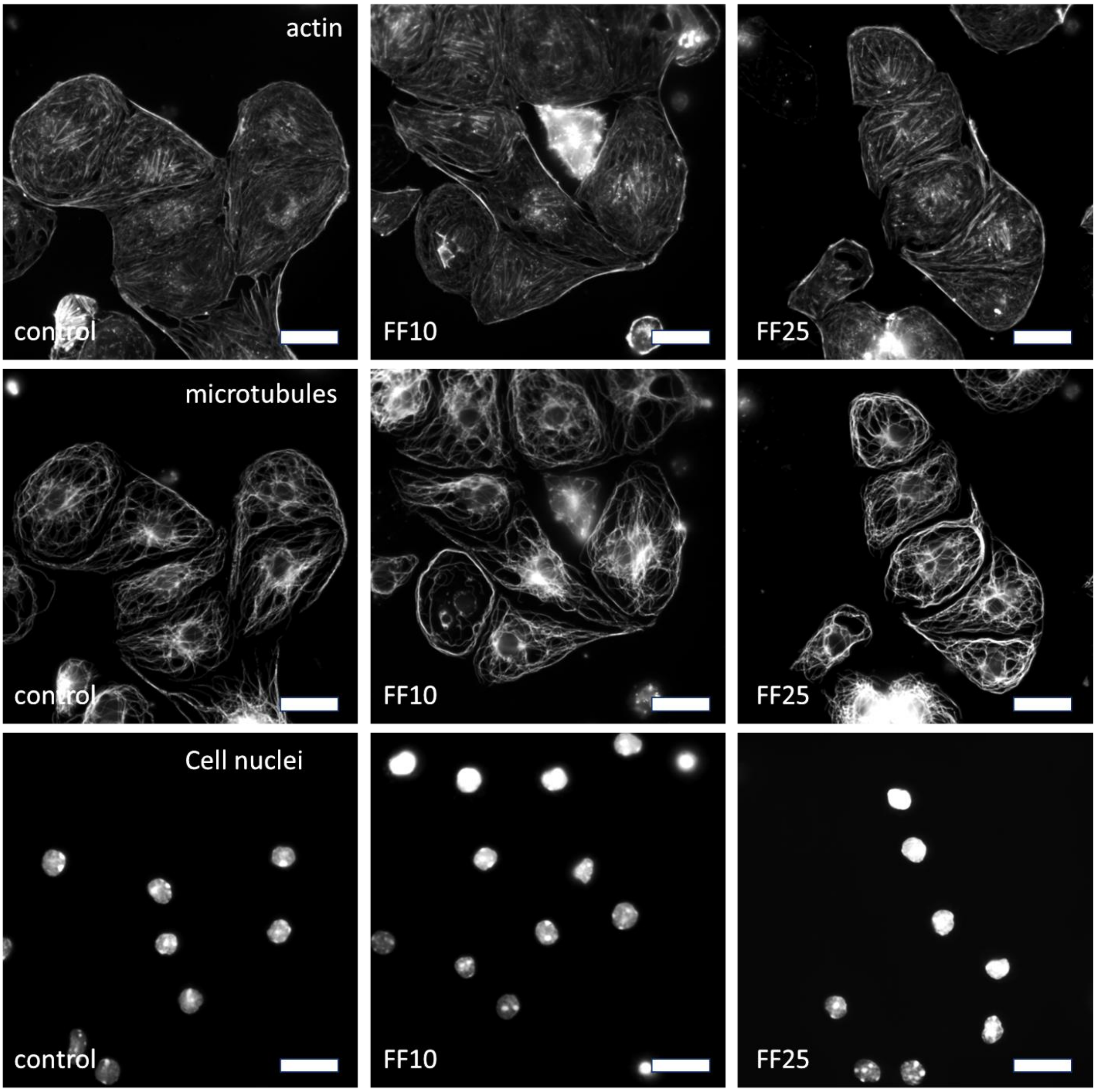


**Supplementary information 3**. **Effect of fenofibrate on changes in the cytoskeleton of LSEC cell**. A) visualization of actin filaments and microtubules and cell nuclei bar represents 25 μm

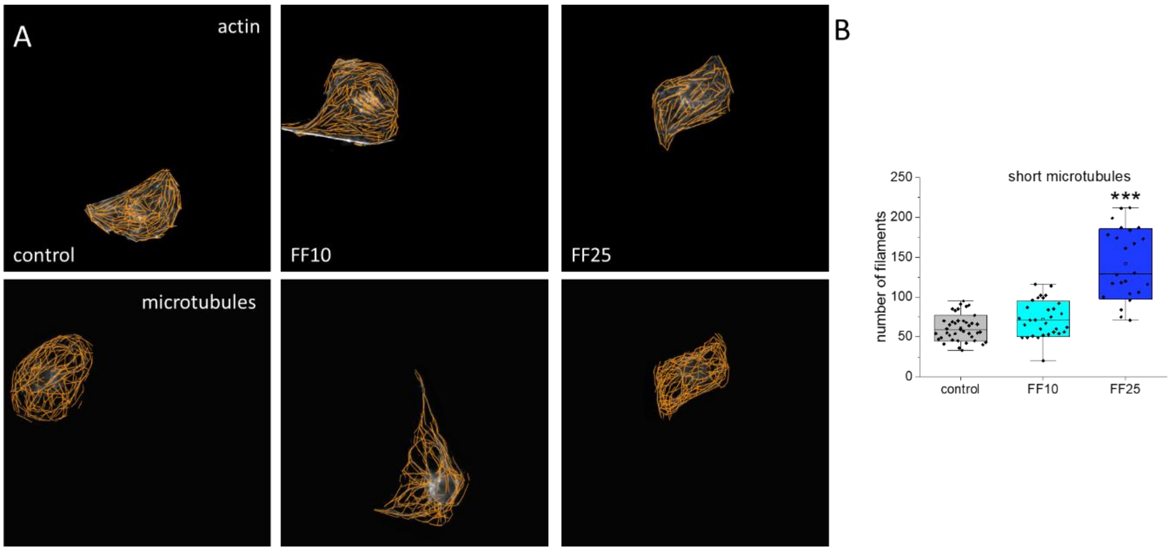


**Supplementary information 4**. **Effect of fenofibrate on changes in the cytoskeleton of LSEC**. A) visualization of actin filaments and microtubules obtained with Filament Sensor software; B) number of short microtubules in cells (3.5-5 um). *Boxes represent standard deviation; dots represent individual cells. Statistical significance (Kruskal-Wallis test with the post-hoc Dunns test): * $p < 0.05$, ** $p < 0.01$, *** $p < 0.001$.*

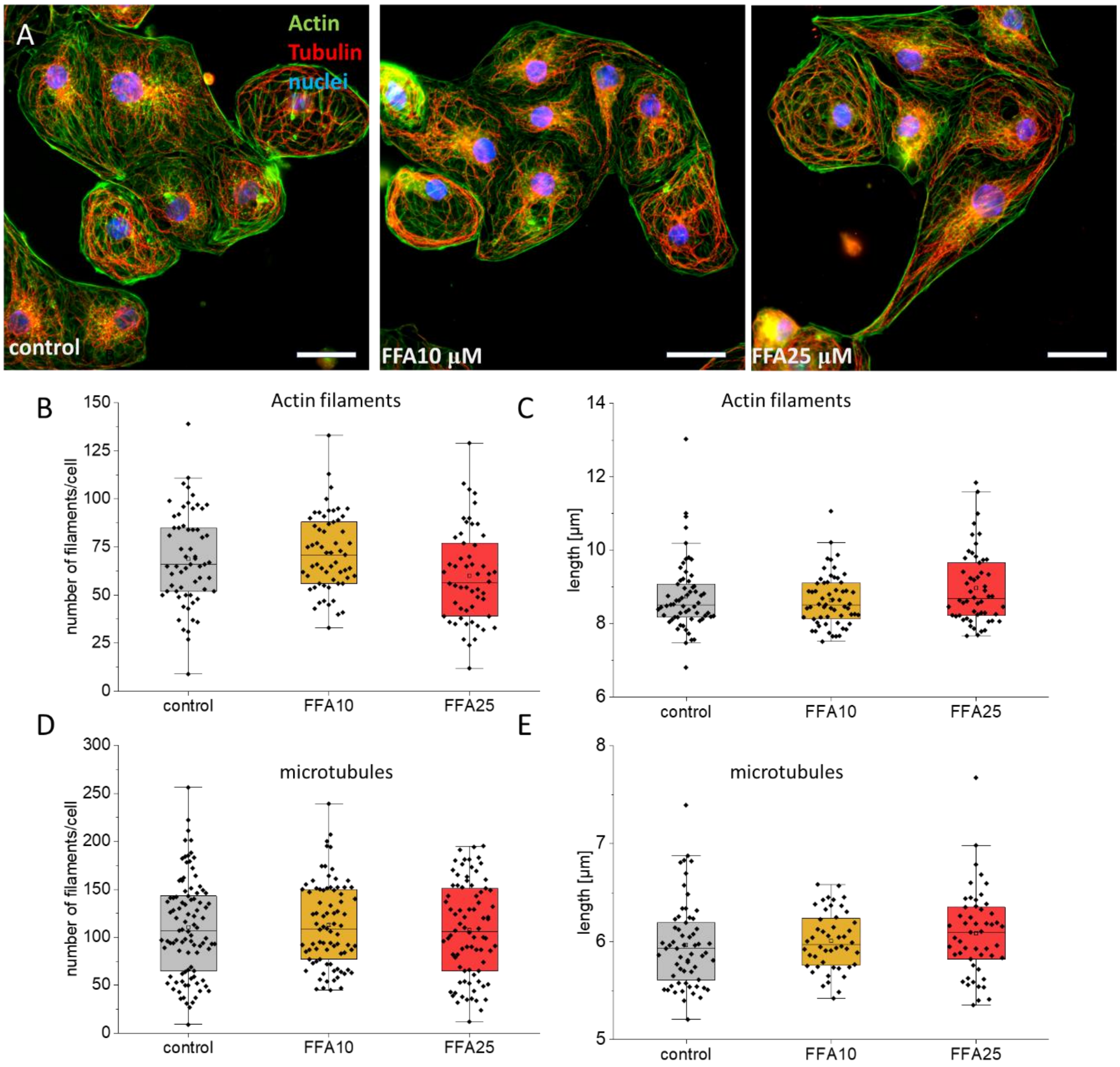


**Supplementary information 5. Effect of fenofibric acid on cytoskeleton of LSEC** . A) visualization of the cytoskeleton of LSEC : (green) actin filaments, (red) microtubules, (blue) cell nuclei, bar presents 25 μm; B) number of actin filaments per cell *(for filaments longer than 5 um*); C) average length of actin filaments in a cell; (*for filaments longer than 5 um*; *dots are presenting separated filaments)*; C) number of microtubules per cell *(for filaments longer than 5 um*); D) average length of microtubules in a cell (*for filaments longer than 5 um)*; *Boxes in B–E represents  SD; dots refers to individual cells; square symbol indicates median and whiskers are 5th and 95th percentiles of the data. Statistical significance (Kruskal-Wallis test with the post-hoc Dunns test): * p < 0.05, ** p < 0.01, *** p < 0.001; n = 4*

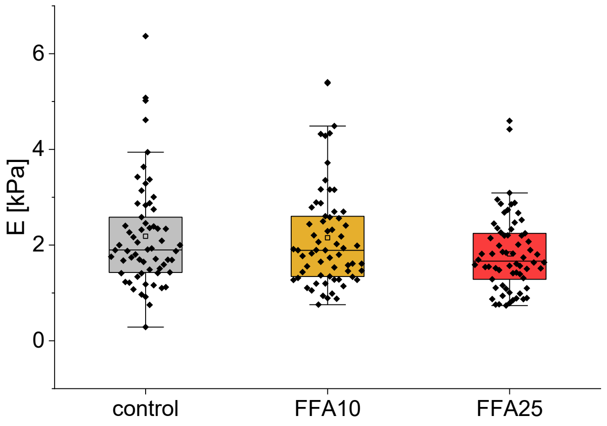


**Supplementary information 6**. **Effect of fenofibric acid on elascisity of LSEC. Young**'s modulus of LSEC measured with AFM *Boxes represent SD; dots refer to individual cells; square symbol indicates median and whiskers are 5th and 95th percentiles of the data. Statistical significance (Kruskal-Wallis test with the post-hoc Dunns testXX model): * p < 0.05, ** p < 0.01, *** p < 0.001; n = 4?.*
